# Ultra-long stable biomimetic nanoparticle Click-ed-to-cancer membrane for anti-cancer treatment

**DOI:** 10.64898/2026.04.19.719453

**Authors:** Rajdeep Chakraborty, Ria Shah, Masuma Akter, Mohammad-Ali Shahbazi, Anastasiia Tukova, Kerwin Shannon

## Abstract

Cancer cell membrane coated biomimetic nanoparticles have been shown to be highly efficient in cellular uptake, homotypic tumour targeting, and the ability to suppress tumour growth compared to uncoated nanoparticles. Long duration anti-cancer treatment regimens require highly stable cancer cell membrane coated biomimetic nanoparticle. To manufacture such highly stable cancer cell membrane coated biomimetic nanoparticle, we used “Click-chemistry” to encapsulate cancer cell membrane on nanoparticles. In situ characterization was done to confirm the functionality of the novel Click-chemistry based formulation to encapsulate cancer cell membrane on nanoparticles. Gold nanoparticles were encapsulated with the cell membranes of cell lines of lung adenocarcinoma, malignant melanoma, high-grade serous epithelial ovarian cancer, colorectal cancer, oral cancer, esophageal adenocarcinoma, adenoid cystic carcinoma of salivary gland, and breast cancer. Functional group analysis, size, morphology, and surface charge confirmed long-stability of the biomimetic nanoparticles after incubating in complete growth medium for 12-months.

## Main

The patients receiving palliative-intent chemotherapy, 30-day mortality directly attributable to chemotherapy was found to be 6.6% [1]. High-risk sub-groups – older patients or those with advanced heavily treated disease (3^rd^ line or higher) experienced significant higher toxicity, with up to 12.6% mortality within 30 days [2]. Research highlights that use of chemotherapy within 30 days of death often results in more adverse events contributing to mortality compared to patients not treated at the end of life. Nanoparticle-based chemotherapy significantly improved overall survival compared to conventional chemotherapy. Significant reduction of hematological toxicity and non-hematological toxicity upon treating the patients with Nanoparticle-based chemotherapy [3]. Consequently, there is a need for new approaches to extend nanoparticle residence time in vivo through particle surface modifications to circumvent macrophage uptake and systemic clearance.

Cancer cell membrane coated biomimetic nanoparticles have been shown to be 20-fold more efficient in cellular uptake compared to uncoated nanoparticles, and 40-fold more efficient than other traditional coating methods [4]. Additionally, Cancer cell membrane coated biomimetic nanoparticles, utilizing membrane proteins, such as E-cadherin, N-cadherin, and galectin-3, show a 6.5-fold increase in homotypic tumour targeting [5]. Biomimetic nanoparticles have demonstrated the ability to suppress tumour growth by around 70%, reduce tumour volume by 60%, and extend survival by 50% in preclinical models [6]. Further, biomimetic nanoparticles commonly achieve high encapsulation efficiency, with some studies reporting approximately 64% efficiency for drugs like Doxorubicin and cisplatin [7]. Cancer cell membrane coated biomimetic nanoparticles allow for a “top-down” approach (creating large amounts of membrane material from cultured cells) compared to the “bottom-up” natural secretion of extracellular vesicles, which typically results in for lower yields [8].

If the anti-cancer treatment duration is long, patients require a cancer cell membrane coated biomimetic nanoparticle with extended shelf life and highly stable in vivo. To manufacture such highly stable cancer cell membrane coated biomimetic nanoparticle, we used “Click-chemistry” to encapsulate cancer cell membrane on nanoparticles.

Gold nanoparticles was encapsulated by tumour membrane using alkyne-PEG-thiol and Azido-PEG-NHS ester, that results in triazole (C_2_H_3_N_3_) linked- to membrane protein. The 1,2,3-triazole moiety is highly stable and does not readily degrade in vivo, making it an ideal, non-toxic linker for drug development [9]. FDA approved Fluconazole for systemic fungal infections are examples of drug using triazole linkage, that are well-tolerated with low toxicity to human cells compared to other anti-fungal drugs [10].

Thiol groups (-SH) form highly stable, covalent, or dative bonds with gold surfaces, commonly used to create self-assembled monolayers (SAMs) for nanotechnology, biosensing, and surface functionalization [11]. The sulfur atom binds to gold, often releasing hydrogen and forming an (Au–S) bond, which is crucial for stabilizing gold nanoparticles and modifying electrical properties. Azide [N=N^+^=N^−^] was linked to alkyne [C_n_H_2n-2_]), forming a triazole that with NHS (N-hydroxysuccinimide) ester group reacted specifically with primary amines, found on lysine residues or the N-terminus of proteins, to form a stable amide bond (CO-NH_2_).

PEG is considered non-toxic and safe for systemic administration (intravenously or subcutaneously) at low-to-moderate molecular doses [12]. Here, our biomimetic nanoparticle linker formulation falls under the classification of low-molecular weight polyethylene glycols. Average polydispersity index of our biomimetic nanoparticle is 0.303 (SD 0.031) and for control gold is 0.110 (SD 0.08) over a period of 12-months (Figure 1). A PDI under 0.3 is commonly used as a quality control criterion indicating that the particles are stable and not severely aggregated. Our biomimetic nanoparticles showed an average zeta potential of approximately −25 mV over a period of 12-months. That further, confirms colloidal stability of the novel biomimetic nanoparticles.

**Figure 1.**
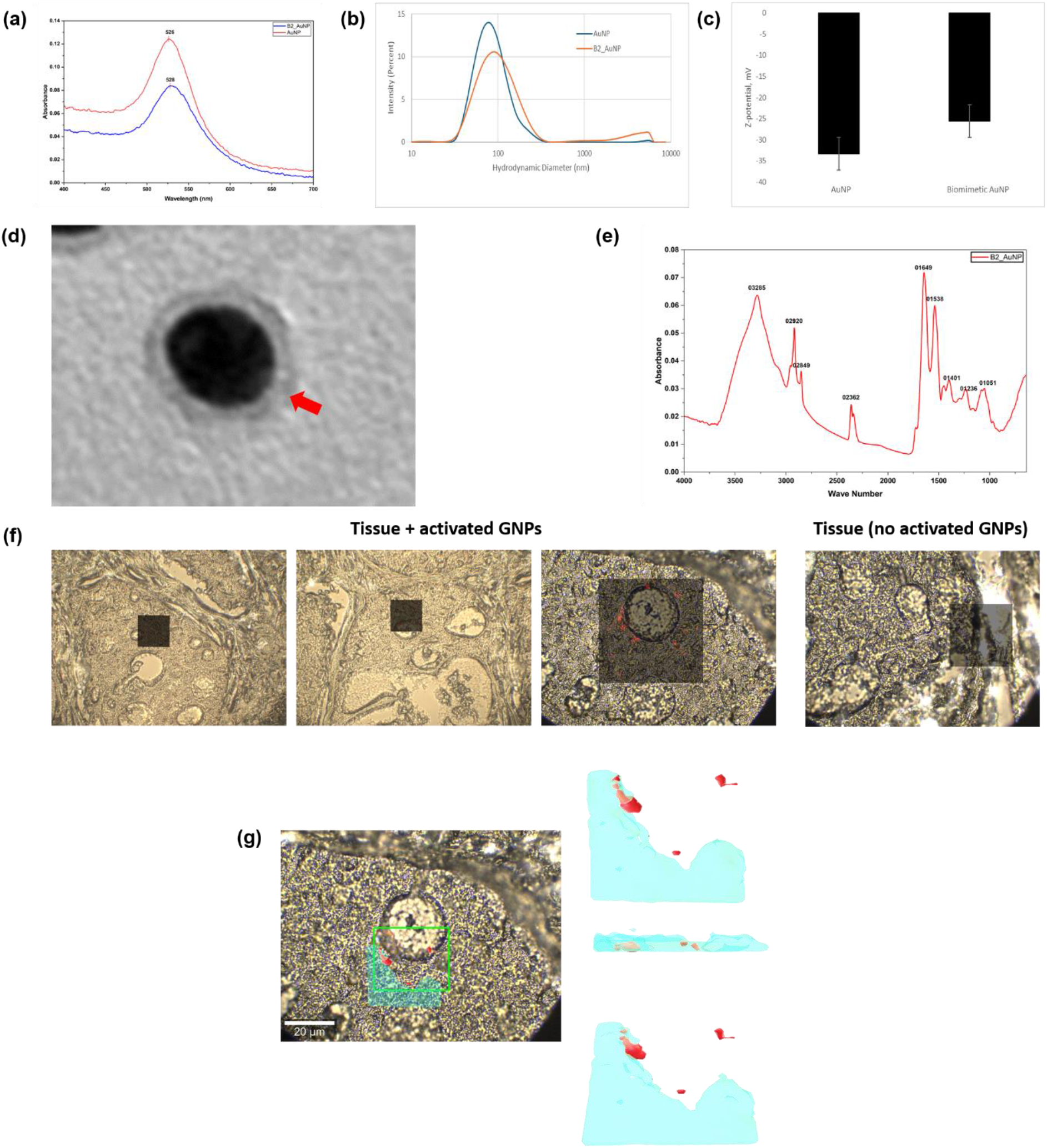
Biophysical chemistry and in-situ characterization of biomimetic gold nanoparticles. (a) UV-Vis spectrum of the biomimetic gold nanoparticles; (b) DLS; (c) Z-potential; (d) TEM of the biomimetic gold nanoparticles showing protein corona around nanoparticles (red arrow); (e) FT-IR of the biomimetic gold nanoparticles; (f) Raman 2D mapping of the biomimetic gold nanoparticles on FFPE slides of adenoid cystic carcinoma of salivary gland (NBP2-30299, Novus Biologicals), showing comparison of Raman signal intensity of sample (Click chemistry activated gold nanoparticles) and control (bare gold nanoparticles). Scale 100 µm. (g) Raman 3D mapping (30 x 25 x 4 µm), red dots indicate regions of high Raman intensity at ~1010 cm^−1^, blue area indicates weak Raman signal form the tissue in 3D Raman mapping. Scale 20 µm.

Using FT-IR, peptide linkage identified – Amide I 1649 cm^−1^ sharp peak confirms conjugation. The sharp peak is primarily due to C=O with minor C-N stretching. 3285 cm^−1^ broad peak denotes O-H stretching and N-H stretching characteristic of amide bonds (-NH), hydroxyl groups (-OH), amino groups (-NH_2_) present in the amino acid backbone and side chains of the protein. This broad peak after gold-protein conjugation, is primarily attributed to the presence of protein layers on the surface of gold nanoparticles. FTIR showing sharp peak 2920 cm^−1^ is characteristic of C-H stretching vibrations that confirms protein attached to surface of gold nanoparticles. Sharp peak at 2362 cm-1 denotes successful covalent bonding reaction (protein linked to a spacer on the nanoparticle). Peaks at 1051 cm-1 denotes presence of capping agents-organic capping shell prevents gold nanoparticles from aggregation (Figure 1).

Upon Raman 2D mapping, red dots indicate regions of high Raman intensity at ~1010 cm^−1^, assigned to the phenyl ring (Phe). This highlights areas where the tissue interacts with the plasmonic surface of the AuNPs. In contrast, tissue regions not in contact with AuNPs do not exhibit measurable Raman signal intensity. Upon Raman 3D mapping, blue area indicates weak Raman signal form the tissue. Baseline signal was treated as absence of Raman signal from tissue or AuNPs (Figure 1).

Upon sub-cellular fractionation, membranes of (i) H1573 cells (lung adenocarcinoma) encapsulated gold nanoparticles incubated in RPMI 1640 Medium (R8758-500 ML), 10% FBS (A56695-01, gibco), 1% Penicillin/Streptomycin solution (G255, abm); (ii) OVCA432 cells (high-grade serous epithelial ovarian cancer) encapsulated gold nanoparticles incubated in Medium 199 (M4530-500ML, Merc Life Science Pty Ltd), MCDB 105 Medium (M6395-1L, Merc Life Science Pty Ltd), 1% Penicillin/Streptomycin solution (G255, abm); (iii) CaCO2 [HTB-37, ATCC] (Colorectal Cancer) encapsulated gold nanoparticles incubated in High-glucose Dulbecco’s modification of Eagle medium (DMEM) (D5796-500ML, Thermo Fisher Scientific), FBS (A56695-01, gibco); (iv) UM-HACC-2A cells (adenoid cystic carcinoma) (T8326, abm) encapsulated gold nanoparticles incubated in optimized salivary gland medium consisted of PriGroIII (TM003, abm), 10% fetal bovine serum (A56695-01, gibco), 2 mM L-glutamine (G275, abm), 0.4 µg/mL hydrocortisone (H0135-1MG, Sigma Aldrich), 20 ng/mL recombinant human epidermal growth factor (Z100139. abm), 5 µg/mL recombinant human insulin (Z101065, abm), 1% Penicillin/Streptomycin solution (G255, abm); (v) A-375 cells (malignant melanoma) (CRL-1619, ATCC) encapsulated gold nanoparticles incubated in High-glucose Dulbecco’s modification of Eagle medium (DMEM) (D5796-500ML, Thermo Fisher Scientific), FBS (A56695-01, gibco), 1% Penicillin/Streptomycin solution (G255, abm); (vi) SCC9 cells (oral cancer) encapsulated gold nanoparticles incubated in High-glucose Dulbecco’s modification of Eagle medium (DMEM) (D5796-500ML, Thermo Fisher Scientific), FBS (A56695-01, gibco), 1% Penicillin/Streptomycin solution (G255, abm); (vii) CW1474 cell line (CRL-3529, ATCC) (esophageal adenocarcinoma) encapsulated gold nanoparticles incubated in High-glucose Dulbecco’s modification of Eagle medium (DMEM) (D5796-500ML, Thermo Fisher Scientific), FBS (A56695-01, gibco), 1% Penicillin/Streptomycin solution (G255, abm); and (viii) MCF7 [HTB-22, ATCC] (Breast Cancer) encapsulated gold nanoparticles incubated in High-glucose Dulbecco’s modification of Eagle medium (DMEM) (D5796-500ML, Thermo Fisher Scientific), FBS (A56695-01, gibco), 1% Penicillin/Streptomycin solution (G255, abm); at 4 °C for 12-months. TEM images showed intact membrane protein corona morphology around gold nanoparticles (Figure 2). Overall, biophysical chemistry, in-situ, and in vitro characterization confirmed the stability of the biomimetic nanoparticles.

**Figure 2.**
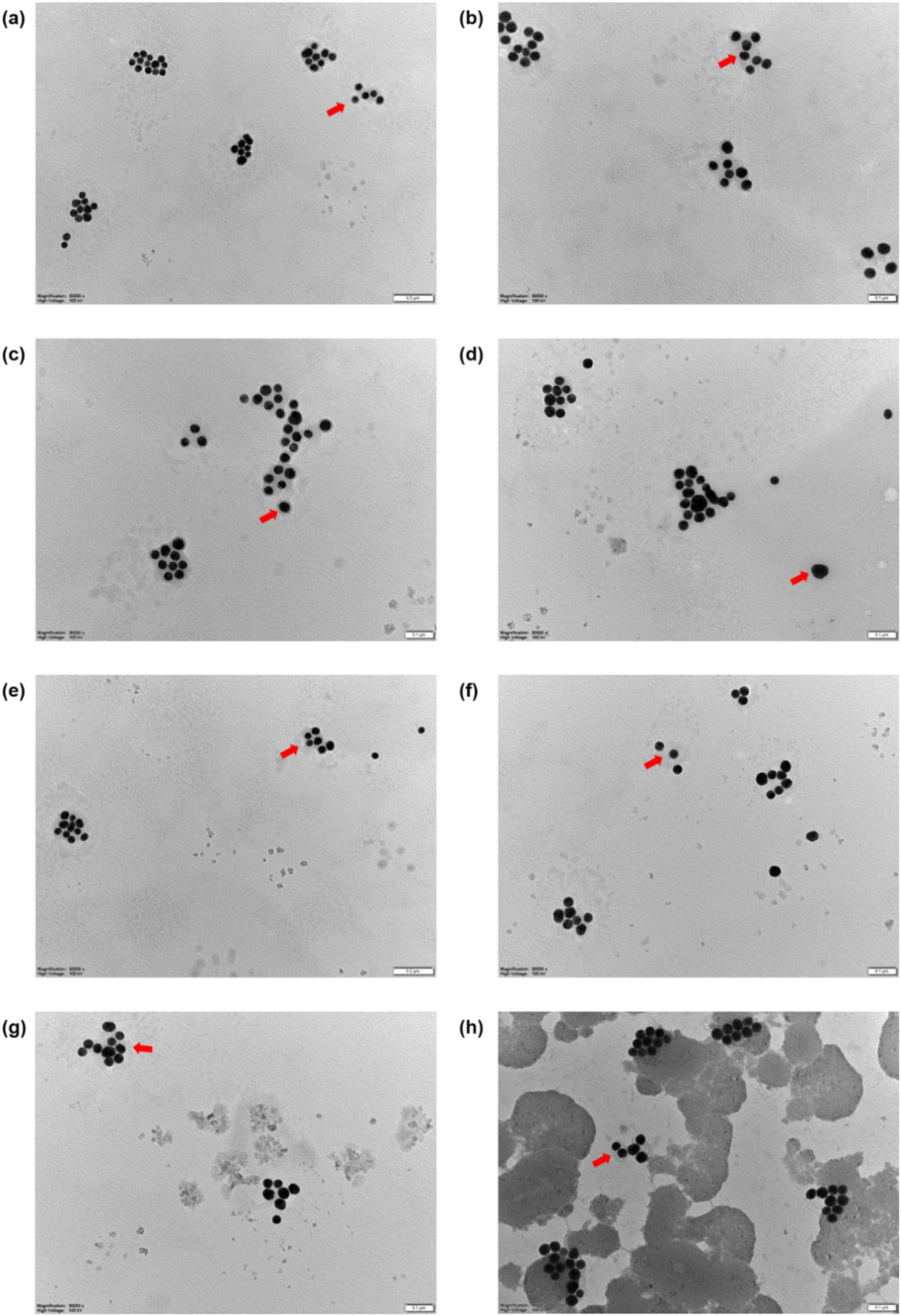
TEM morphological images of biomimetic nanoparticles after 12-months incubation. Red arrow indicates membrane protein corona around the biomimetic gold nanoparticles of (a) lung adenocarcinoma, (b) malignant melanoma, (c) high-grade serous epithelial ovarian cancer, (d) colorectal cancer, (e) oral cancer, (f) esophageal adenocarcinoma, (g) adenoid cystic carcinoma of salivary gland, and (h) breast cancer, at magnification 80,000X, Scale 0.1 µm.

## Conclusion

International Council for Harmonisation (ICH) guidelines for drug products typically requires long-term stability study for at least 12-months, making the duration the gold standard for long-term storage. The spacer arm length of a PEG12 (12-unit polyethylene glycol) spacer used in the construction of our biomimetic nanoparticle is approximately 46.8 Å to 56.0 Å (4.68 to 5.6 nm). PEG12 provides long-duration stability to nanoparticles by creating a sterically hindered, hydrophilic, and neutral surface layer that prevents aggregation and resists interaction with biological components. PEG12 serves as a functional stealth coating by acting as a non-ionic hydrophilic polymer, enhancing both physical shelf-life and in vivo circulation time [13]. Future in vivo effect based on the homing effect of this novel biomimetic nanoparticle formulation should be determined using FDA approved PLGA nanoparticles. [14]

## Methods

### (1) Biomimetic nanoparticle preparation

H1573 cells (lung adenocarcinoma) (CRL-5877, ATCC) cultured in RPMI 1640 Medium (R8758-500 ML), 10% FBS (A56695-01, gibco), 1% Penicillin/Streptomycin solution (G255, abm); OVCA432 cells (high-grade serous epithelial ovarian cancer) (cells kindly gifted by Dr Charlie Ahn, Macquarie University) cultured in Medium 199 (M4530-500ML, Merc Life Science Pty Ltd), MCDB 105 Medium (M6395-1L, Merc Life Science Pty Ltd), 1% Penicillin/Streptomycin solution (G255, abm); CaCO2 [HTB-37, ATCC] (Colorectal Cancer) cultured in High-glucose Dulbecco’s modification of Eagle medium (DMEM) (D5796-500ML, Thermo Fisher Scientific), FBS (A56695-01, gibco); UM-HACC-2A cells (adenoid cystic carcinoma) (T8326, abm) cultured in optimized salivary gland medium consisted of PriGroIII (TM003, abm), 10% fetal bovine serum (A56695-01, gibco), 2 mM L-glutamine (G275, abm), 0.4 µg/mL hydrocortisone (H0135-1MG, Sigma Aldrich), 20 ng/mL recombinant human epidermal growth factor (Z100139. abm), 5 µg/mL recombinant human insulin (Z101065, abm), 1% Penicillin/Streptomycin solution (G255, abm); A-375 cells (malignant melanoma) (CRL-1619, ATCC) cultured in High-glucose Dulbecco’s modification of Eagle medium (DMEM) (D5796-500ML, Thermo Fisher Scientific), FBS (A56695-01, gibco), 1% Penicillin/Streptomycin solution (G255, abm); SCC9 cells (oral cancer) (cells kindly gifted by Dr Charbel Darido, University of Melbourne) cultured in High-glucose Dulbecco’s modification of Eagle medium (DMEM) (D5796-500ML, Thermo Fisher Scientific), FBS (A56695-01, gibco), 1% Penicillin/Streptomycin solution (G255, abm); CW1474 cell line (CRL-3529, ATCC) (esophageal adenocarcinoma) cultured in High-glucose Dulbecco’s modification of Eagle medium (DMEM) (D5796-500ML, Thermo Fisher Scientific), FBS (A56695-01, gibco), 1% Penicillin/Streptomycin solution (G255, abm); and MCF7 [HTB-22, ATCC] (Breast Cancer) cultured in High-glucose Dulbecco’s modification of Eagle medium (DMEM) (D5796-500ML, Thermo Fisher Scientific), FBS (A56695-01, gibco), 1% Penicillin/Streptomycin solution (G255, abm). Mem-PER Plus Membrane Protein Extraction Kit (89842, Thermo Fisher Scientific) was used to isolate membrane protein of the cancer cells using the manufacturer’s protocol of sub-cellular fractionation. Gold (Au) nanoparticles of 40 nm were used during the project. In-house developed Click chemistry protocol was used to encapsulate the cancer cell membrane on gold nanoparticles (Novel biomimetic nanoparticle, Patent Number 2026901133, IP Australia). Upon encapsulation of the cancer cell membrane on the nanoparticles, the nanoparticles were incubated in the cancer cell corresponding full growth culture medium at 4 °C for 12-months.

### (2) Biophysical Chemistry Characterization

Optical properties of biomimetic gold nanoparticles were assessed using UV-visible spectroscopy. Spectra were acquired over a wavelength range of 200-900 nm using JASCO V-760 UV-vis spectrometer (Jasco, Hachioji Japan). The measurements were acquired using 1000 µL of the sample placed in a plastic disposable semi-micro cuvette (Brand Tech, USA). The surface charge and size (dynamic light scattering) of the nanoparticles were determined using Zetasizer ZS (Malvern Panalytica, Malvern, U.K). Assessment of amide linkage to determine cancer cell membrane encapsulation of nanoparticles were done using iN10 FT-IR microscope (Thermo Fisher Scientific).

### (3) FFPE sample processing

Two salivary gland condition FFPE slides were used during the project. (i) Adenoid cystic carcinoma of salivary gland (NBP2-30299, Novus Biologicals), Layout: 1X1, Diameter: 4, Thickness: 5 µm, Age: 51 years, Sex: F, Organ: Parotid gland, Pathology: Adenoid Cystic Carcinoma, Tissue Status: Tumor/Cancer, Species: Human; (ii) normal parotid salivary gland (NBP2-30192, Novus Biologicals), Layout: 1X1, Diameter: 4, Thickness: 5 µm, Age: 37 years, Sex: F, Organ: Parotid gland, Tissue Status: Normal, Species: Human. Deparaffinization process initiated with FFPE samples kept in 60 °C in oven for 1 hour, followed by immersing in xylene two-times for 10 minutes in each round, then 100% ethyl alcohol for 10 minutes, then 95% ethyl alcohol for 10 minutes, then 70% ethyl alcohol for 10 minutes, and finally rehydrated in distill water for 5 minutes.

### (4) In situ characterization

For Raman 2D mapping, the area 30 x 30 µm was scanned with 633 nm laser with 20 mW power with 1 second exposure under 50X lens. In-plane resolution is about 0.6 µm/pixel in both directions. For 3D Raman mapping, area 30 x 25 x 4 µm was scanned with 633 nm laser with 20 mW power with 1 s exposure under 50X lense. With 0.6 × 0.5 µm in-plane resolution, with ~1 µm axial spacing between z-slices.

### (5) Morphology assay

The morphology of biomimetic gold nanoparticles was assessed after acquiring transmission electron microscopy images (JEOL 1400 TEM, Tokyo, Japan).

## Author contributions

Conceptualization: RC; Project and experimental design: RC, MS, AT, KS; Funding Acquisition: RC, KS; Methods: RC, RS, MS, AT, KS; Data acquisition: RC, RS, MA, AT; Statistical Analysis: MS, AT; Manuscript write up: RC; Manuscript corrections: RS, AT, MS, KS; Supervision: KS

## Ethics

The project was done after Macquarie University Medical Sciences Human Research Ethics Committee approval to use FFPE samples “19710 - Chakraborty-A Prelude towards transforming adenoid cystic carcinoma into an immune hot tumour”. Cell and spheroid immune coculture work was done upon acquiring biosafety approval from Institutional Biosafety Committee, Macquarie University, “A tumour-cell based initiation of immune evasion – 16835”.

## Acknowledgement

The project is funded by the Adenoid Cystic Carcinoma Research Foundation (ACCRF, Massachusetts, USA) and Australian and New Zealand Head and Neck Cancer Society (ANZHNCS, Australia). We are grateful to Jeffrey Kauffman (Co-founder of ACCRF) and Dr Nicole Spardy Burr (Executive Director, ACCRF) for their immense help towards the project and providing me useful information and suggestions regarding experimental design of the project. We are grateful to the scientific committee of ANZHNCS for their continuous trust and support towards innovative cancer research. We would like to acknowledge the support from Prof Yuling Wang group at the School of Natural Sciences, Macquarie University, and the staff of microscopy unit and Australian Proteome Analysis Facility, Faculty of Science and Engineering, Macquarie Analytical and Fabrication Facility (MAFF). Macquarie University. We would like to acknowledge the support of Prof Jacques Nor (University of Michigan), Prof Fredrick Kaye (University of Florida), Prof Fiona Simpson (University of Queensland), and Dr Charbel Darido (University of Melbourne) for their initial technical support during the culturing of adenoid cystic carcinoma cells.

